# Adenine DNA methylation associated to transcription is widespread across eukaryotes

**DOI:** 10.1101/2024.10.28.620566

**Authors:** Pedro Romero Charria, Cristina Navarrete, Vladimir Ovchinnikov, Luke A Sarre, Victoria Shabardina, Elena Casacuberta, David Lara-Astiaso, Arnau Sebé-Pedrós, Alex de Mendoza

## Abstract

DNA methylation in the form of 5-methylcytosine (5mC) is widespread in eukaryotes, while the presence of N6-methyladenine (6mA) has sparked considerable debate. Methodological disparities in quantifying and mapping 6mA in genomic DNA have fueled this controversy. Yet, the distantly related early branching fungi, ciliates and the algae *Chlamydomonas reinhardtii* exhibit robust 6mA methylation patterns, but the origin and evolution of 6mA remain unknown. Here we use Oxford Nanopore modified base calling to profile 6mA at base pair resolution in 18 unicellular eukaryotes spanning all major eukaryotic supergroups. Our results reveal that only species encoding the adenine methyltransferase AMT1 display robust genomic 6mA patterns. Notably, 6mA consistently accumulates downstream of transcriptional start sites, aligning with H3K4me3-enriched nucleosomes, suggesting a conserved role in placing transcriptionally permissive nucleosomes. Intriguingly, the recurrent loss of the 6mA pathway across eukaryotes, particularly in major multicellular lineages, implies a convergent alteration in the dual methylation system of the Last Eukaryotic Common Ancestor, which featured transcription-associated 6mA and repression-associated 5mC.

## Main text

Chemical modifications can impact the functions of the four main DNA nucleobases. The most studied modification is the addition of a methyl group to the fifth carbon ring of cytosines, 5mC. In eukaryotes, 5mC is a widespread form of DNA methylation deposited by the family of DNA methyltransferase enzymes (DNMTs)^1–3^, and its functions range from transposable element control to gene regulation^4–6^. Other types of base modifications, like hydroxymethyluracil in dinoflagellates or the “base J” in kinetoplastids, are lineage-restricted but abundant in some species^7,8^. While 5mC is studied as the major eukaryotic DNA methylation form, 6-methyladenine (6mA) has been reported across various lineages, including animals^9–12^, plants^13^, early diverging fungi^14^, algae^15^, and ciliates^16,17^. However, global 6mA levels are often close to detection limits, with reports questioning its presence in animals, plants, or yeast^18–22^. Challenges in 6mA quantification include antibody pull-down specificity issues, bacterial DNA contamination, RNA contamination, and sequencing data artefacts^23,24^. Despite this, ciliates, early diverging fungi, and the algae *Chlamydomonas reinhardtii* exhibit relatively high and orthogonally confirmed 6mA levels^14,18^, highlighting the presence of this modification in some eukaryotes. Inadequate taxon sampling and contradictory technical results have therefore resulted in a limited understanding of the distribution and evolution of 6mA.

A mechanistic understanding of genomic 6mA in eukaryotes comes from research in ciliates. In ciliates, the active methyltransferase enzymes, belonging to the MT-A70 family, differ from DNMTs but share structural homology with well-characterised 6mA RNA methyltransferases, METTL3 and METTL14^25–27^. The crucial writer of genomic 6mA in ciliates is AMT1 (known as MTA1 in *Oxytricha trifallax*), while accessory enzymes from families AMT6/7 (MTA9 in *O. trifallax*) are dispensable and lack catalytic activity^17,28^. However, AMT6/7 forms a heterodimer with AMT1 during DNA engagement, akin to the METTL3 and METTL14 heterodimer in RNA methylation (**Fig. 1a**)^29^. Additionally, DNA-binding proteins p1 and p2, originally described in the ciliate *O. trifallax,* play a role in targeting genomic DNA by forming a multimeric complex with AMT1 ^17,28^. In ciliates, early diverging fungi, and the algae *C. reinhardtii*, 6mA is confined to ApT dinucleotides^14–17,27^. It is suggested that AMT1 binds hemimethylated ApT sites, similar to the role of DNMT1 in maintaining 5mC on CpG dinucleotides across cell divisions^30^. In ciliates and *C. reinhardtii*, 6mA is found in internucleosomal linker DNA^15–17,31^, particularly enriched downstream of the transcriptional start site (TSS) of active genes. This association with transcription contrasts the proposed role of 6mA in silencing transposable elements in animals^11,12,21^. 6mA may be an evolutionary flexible modification as observed for 5mC, which has evolved roles exclusively in transposon silencing or transcribed gene bodies in different eukaryotes^4–6^. However, the lack of comprehensive data from several branches of the tree of life hampers our ability to reconstruct 6mA evolution and functions.

**Fig. 1.**
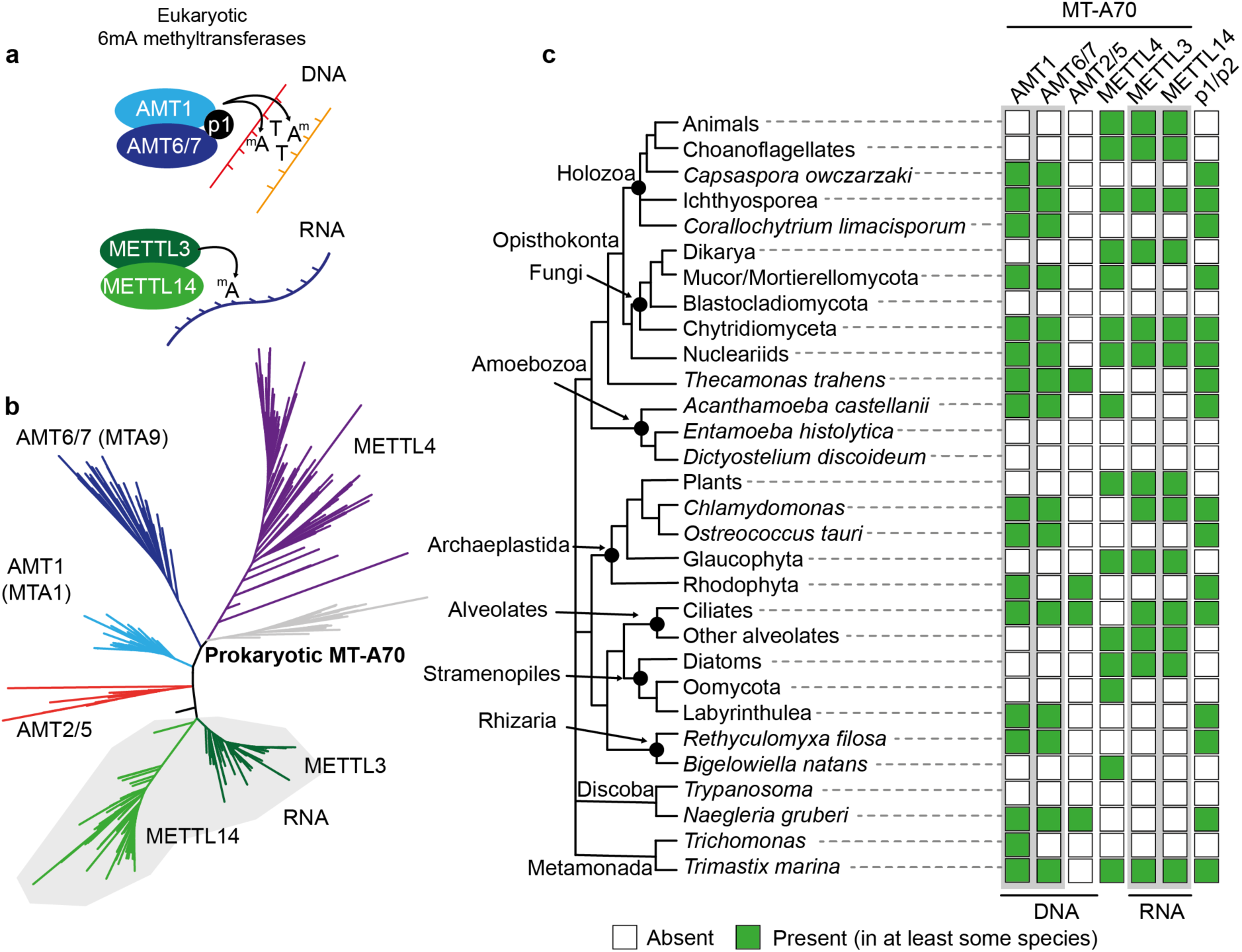
MT-A70 6-methyl-adenine methyltransferases origin and recurrent loss. (**a**) Diagram of the known heterodimers formed by MT-A70 6mA methyltransferases. AMT1 and METTL3 are the active enzymes whereas their partners are required but do not catalyse the reaction. p1 and p2 are described proteins from the ciliate *O. trifallax* that are required for AMT1 function^17^. (**b**) Maximum likelihood phylogenetic tree of MT-A70 6mA methyltransferases, including eukaryotic and prokaryotic sequences. Each clade is named as previous references. (**C**) Distribution of MT-A70 6mA methyltransferase family members across eukaryotes. For some lineages the presence suggests that at least some species within the clade contains them. Phylogenetic relationships across these species are based on consensus in the field^34,92^.

To clarify these uncertainties, in this work we study the evolutionary history of MT- A70 methyltransferases and we find that the AMT1 pathway is ancestral in eukaryotes. Then, utilising Oxford Nanopore modified base-calling, previously validated for identifying this base modification in eukaryotes^32^, we detect consistent 6mA patterns in representatives from the major eukaryotic supergroups, thereby tracing 6mA back to the Last Eukaryotic Common Ancestor (LECA).

## Results

### The AMT1 pathway is ancestral but recurrently lost in eukaryotes

To clarify the origins and evolution of MT-A70 methyltransferases, we examined 231 eukaryotic genomes, incorporating transcriptomes from lineages lacking genome data. Additionally, we used eukaryotic MT-A70 sequences to identify non-eukaryotic homologues, as the origin of these is not well understood^1,25^. The resulting MT-A70 phylogenetic tree mirrors the six established eukaryotic families: AMT1, AMT6/7, AMT2/5, METTL3, METTL14, and METTL4 (**Fig. 1b**)^25,27^. Notably, prokaryotic sequences form a distinct clade, suggesting a bacterial origin for MT-A70 inherited early in eukaryotic evolution (**Fig. 1b, Extended Data Fig. 1a**). MT-A70 presence in prokaryotes is confined to select eubacterial phyla, with only four archaean genomes harbouring it, typically in a single copy per species (**Extended Data Fig. 1b**). Unlike the more widespread bacterial-type Dam 6mA methyltransferase, MunI-like 6mA MT-A70 in bacteria are less diverse^1^. The monophyletic clustering of bacterial sequences, without enrichment in a specific bacterial lineage, indicates a single origin of eukaryotic MT-A70 from an unidentified bacterial donor. This rules out recurrent transfers from prokaryotes to eukaryotes or multiple horizontal transfers from eukaryotes to prokaryotes.

By mapping the distribution of MT-A70 in eukaryotes, we found that all six families are present in most of the eukaryotic supergroups (**Fig. 1c**), suggesting that they are the product of duplication events that occurred before LECA. The RNA-associated METTL3 and METTL14 co-occur across the species phylogeny, in a pattern similar to the DNA-associated AMT1 and AMT6/7, suggesting that both enzymatic pairs have conserved heterodimer associations across eukaryotes, with the loss of one co-occuring with the loss of the other members (**Fig. 1c, Extended Data Fig. 1c**). AMT2/5 is less common, often co-occuring with AMT1, hinting at a potential partnership but one in which AMT2/5 is dispensable. Notably, AMT2/5 is exceptional as it possesses C-terminal ZZ zinc finger domains, yet the function of these is unknown (**Extended Data Fig. 1d**)^25,27^. In contrast, METTL4 does not co-occur with any MT-A70 families, suggesting it does not form heterodimers (**Fig. 1c, Extended Data Fig. 1c**). METTL4 is the only MT-A70 found in fission yeast, which has been shown to have negligible levels of 6mA^19^. In fact, METTL4 is believed to methylate mitochondrial DNA or small RNAs, but its role is currently not well understood^21,33^. Similarly, the nematode *Caenorhabditis elegans* METTL4 orthologue (known as DAMT-1) has been suggested as an active genomic DNA methyltransferase^10^, yet the patterns of 6mA in nematodes are also contested^19^. Beyond the MT-A70 methyltransferases, the proteins p1 and p2 described in *O. trifallax* exclusively co-occur with AMT1 (**Fig. 1c**), indicating an ancestral link with the AMT1 pathway^17^.

Hence, the enzymes responsible for RNA and DNA 6mA exhibit ancient origins, yet their evolution appears largely decoupled to each other in eukaryotes. Importantly, AMT1 is consistently found in species with documented robust 6mA levels, such as ciliates, *C. reinhardtii*, and early fungi^14–17^. AMT1 presence seems reliant on the occurrence of an AMT6/7 and p1/p2 partners, though exceptions exist (e.g. *Trichomonas vaginalis*).

### Genomic 6mA is associated with transcription in AMT1-encoding eukaryotes

Considering the distribution of AMT1 among eukaryotes, we sought to identify lineages exhibiting detectable genomic 6mA. To ensure accuracy and avoid potential bacterial DNA contamination, we focused on species grown axenically under laboratory conditions, and with high-quality genomes available. Our selection included 15 species encoding AMT1 and three species lacking AMT1 orthologues as negative controls. Our sampling of ichthyosporeans, *Corallochytrium limacisporum*, a chytrid fungi, amoebozoans, chlorophytes, a glaucophyte, stramenopiles, heteroloboseans, and a metamonada spans major eukaryotic branching groups^34^. Genomic DNA was sequenced using Oxford Nanopore R9.4.1 technology for 14 species, while equivalent raw Nanopore data from public repositories were used for 4 others^32,35–37^. Employing the Guppy software with the Oxford Nanopore pre-trained “all-context” model, we achieved base pair resolution maps for 6mA and 5mC in all species. Global methylation levels varied significantly across species, with the lowest levels observed in species lacking AMT1 (**Fig. 2a**).

**Fig. 2.**
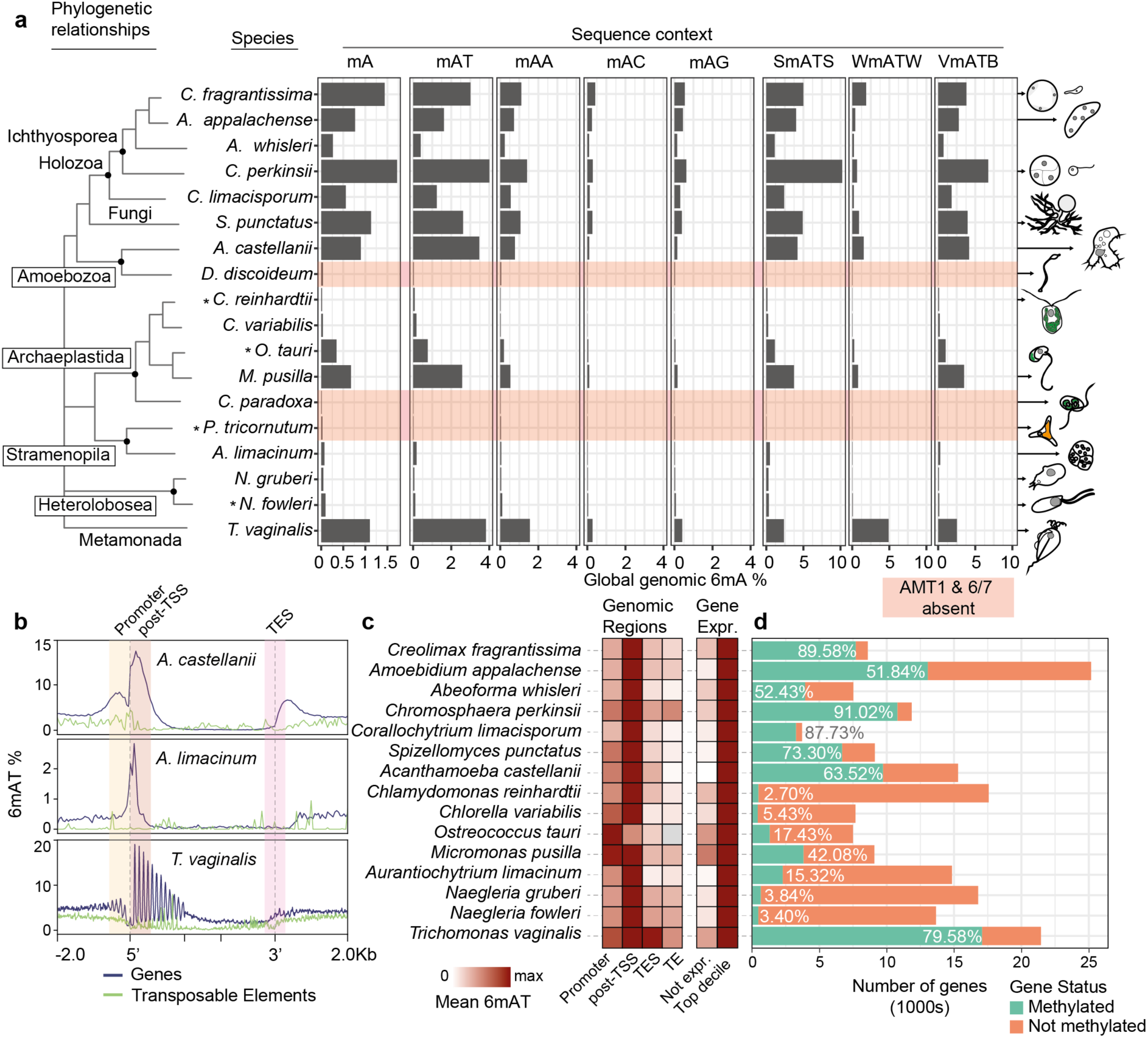
Nanopore sequencing reveals widespread 6mA in AMT1 encoding eukaryotic genomes. (**a**) Global methylation levels of 6mA in different sequence contexts across 18 eukaryotes representing deep branching points of eukaryotic diversity. S stands for C or G, W stands for A or T, V stands for A,C,G and B stands for T,C,G. Species with red shade do not encode for AMT1 or its partner AMT6/7. Asterisks indicate Oxford Nanopore datasets obtained from NCBI/ENA, silhouettes represent the shapes of the species. (**b**) Average 6mA levels in the AT context over 3 example species, including genes (blue) and TEs (green, >1000 bp). 5’ and 3’ ends of the features are marked with dashed lines, first and last 1500 bp are unscaled, and the shades indicate the regions used in panel C. (**c**) Heatmap displaying average 6mA levels across various genomic regions, including the 500 bp downstream of the Transcriptional Start Site (post-TSS), the 500 bp centred around the Transcriptional End Site (TES), and Transposable Elements (TE). Also, gene body 6mA levels for non-expressed genes and top decile of highly expressed genes, see full values in **Extended Data Fig. 3**. Grey cell indicates missing data, as *O. tauri* has too few repeats. (**d**) Number of genes methylated in each genome. A gene is defined as methylated if it contains at least 3 ApTs with >10% 6mA, and coverage above 10x in all species except in *M.pusilla* and *A.whisleri* where 5x was used and *C. limacisporum* and *C.variabilis* where 3x was used.

Given the preference for ApT dinucleotide methylation described for AMT1^14–16^, we assessed global 6mA levels across various sequence contexts. The predominant methylated dinucleotide context in all AMT1-encoding species was AT, though the AA context was not insignificant (**Fig. 2a**). Consistent with findings in ciliates and *C. reinhardtii*, ApT sites displayed symmetric methylation across all AMT1-encoding species (**Extended Data Fig. 2a**). Notably, *C. reinhardtii* is suggested to exhibit higher methylation levels in the VATB context (V = C/G/A, B = C/G/T)^18^. Thus, we calculated global levels for three 4-mer combinations to discern flanking specificity to the AT context. Across all AMT1-containing species, except one, SATS (S = C or G) exhibited the highest methylation, while WATW (W = A or T) showed the lowest (**Fig. 2a**). The VATB context displayed intermediate levels, suggesting that cytosine and guanine flanking might dictate methyltransferase specificity (**Fig. 2a**). An exception to this flanking preference was observed in the metamonad *Trichomonas vaginalis*, which encodes four AMT1 paralogues but lacks AMT6/7 orthologues (**Fig. 1c**). In this species, WATW was more enriched than SATS (**Fig. 2a**), indicating a distinct sequence preference for methyltransferases and highlighting that the observed preferences are not a result of biases in Oxford Nanopore base calling models.

We proceeded to investigate the genomic distribution of 6mA deposition. In species lacking AMT1, we observed uniform 6mA patterns across gene bodies and transposable elements (**Extended Data Fig. 2b**), indicating that the negligible 6mA levels are likely background noise. Conversely, in the remaining species, a distinctive peak of AT methylation emerged proximal to the Transcriptional Start Site (TSS) (**Fig. 2b,c**). Intriguingly, AA methylation mirrored these patterns, suggesting that AA methylation might be an off-target substrate of AMT1 (**Extended Data Fig. 2c**). Some species displayed a subtle 6mA enrichment in promoter regions (upstream of the TSS, **Fig. 2c**)^15^, which, upon scrutiny, proved to be an artefact arising from the head-to-head orientation of upstream genes (**Extended Data Fig. 2d**). Neither Transcriptional End Sites nor Transposable Elements exhibited enrichment in 6mA across any species (**Fig. 2c**). Genic 6mA correlated with gene transcriptional activity (**Fig. 2c**). Across all species, silent genes presented lower 6mA levels compared to highly expressed genes, which also harboured more methylated ApT sites (**Fig. 2c, Extended Data Fig. 3**). Notably, the association with transcription was quite variable; for instance, *Acanthamoeba castellanii* exhibited a gradual link with transcriptional levels, while in others, like ichthyosporeans or prasinophytes, upon a specific transcriptional threshold resulted in uniform 6mA levels across all genes (**Extended Data Fig. 3**). This diversity was also evident in the total number of genes displaying methylation in each genome (**Fig. 2d**). For instance, the ichthyosporean *Chromosphaera perkinsii* exhibited 92% of genes with 6mA, whereas both *Naegleria* species displayed detectable methylation in only 3% of their genes (**Fig. 2d**), showing a widespread variability in amount of 6mA use across eukaryotes.

We then tested how variable 6mA is upon transcriptional changes. We chose the ichthyosporean *Creolimax fragrantissima* and the amoebozoan *A. castellannii* because these have well defined cell stages that can be isolated in the culture^38,39^, high 6mA levels, and distinct associations between 6mA and transcriptional level in the steady state (**Extended Data Fig. 3**). Both species presented highly static 6mA methylomes at base pair and gene level resolution, where most genes have minimal 6mA differences (**Extended Data Fig. 4a,b**). However, in *A. castellanii*, genes that have higher 6mA in the trophozoite stage are also more highly transcribed in this stage, and become silenced during the encystment process. Conversely, genes with higher 6mA in cysts did not show higher transcription, yet this might be a consequence of the cysts being a quiescent cell state (**Extended Data Fig. 4b,c**). In contrast, *C. fragrantissima* amoeba and coenocyte stages presented lower gene level 6mA differences, and the genes that showed more divergent 6mA levels across samples were not transcriptionally different (**Extended Data Fig. 4b,c**). Moreover, differentially transcribed genes among *C. fragrantissima* stages were not among those with different 6mA levels. This is consistent with 6mA having a more gradual and fine tuned link with transcription in *A. castellannii*, while in *C. fragrantissima* all genes within a certain transcriptional threshold showed equivalent 6mA levels. In sum, 6mA is consistently associated with transcription in all AMT1-encoding eukaryotes, yet the dynamic link with transcription and genomic abundance varies across lineages.

### 5mC and 6mA are functionally compartmentalised in eukaryotic genomes

As 5mC represents the other form of DNA methylation widespread in eukaryotes, we investigated its relationship with 6mA within our species set (**Fig. 3a**). We found diverse scenarios: species like *Corallochytrium limacisporum* exhibited non-detectable 5mC levels alongside high 6mA, while others, like the diatom *Phaeodactylum tricornutum*, exclusively harboured 5mC. Some species presented both modifications, and one, *Dictyostelium discoideum*, displayed neither (**Fig. 3a**). Importantly, the negligible 5mC levels detected via Oxford Nanopore aligned with previous findings using alternative techniques, including Enzymatic Methyl-seq or Whole Genome Bisulfite sequencing for some of these same species^40,41^. This variation of patterns suggests that the presence of 6mA is not evolutionarily linked to that of 5mC, corroborating observations in early diverging fungal lineages^14^.

**Fig. 3.**
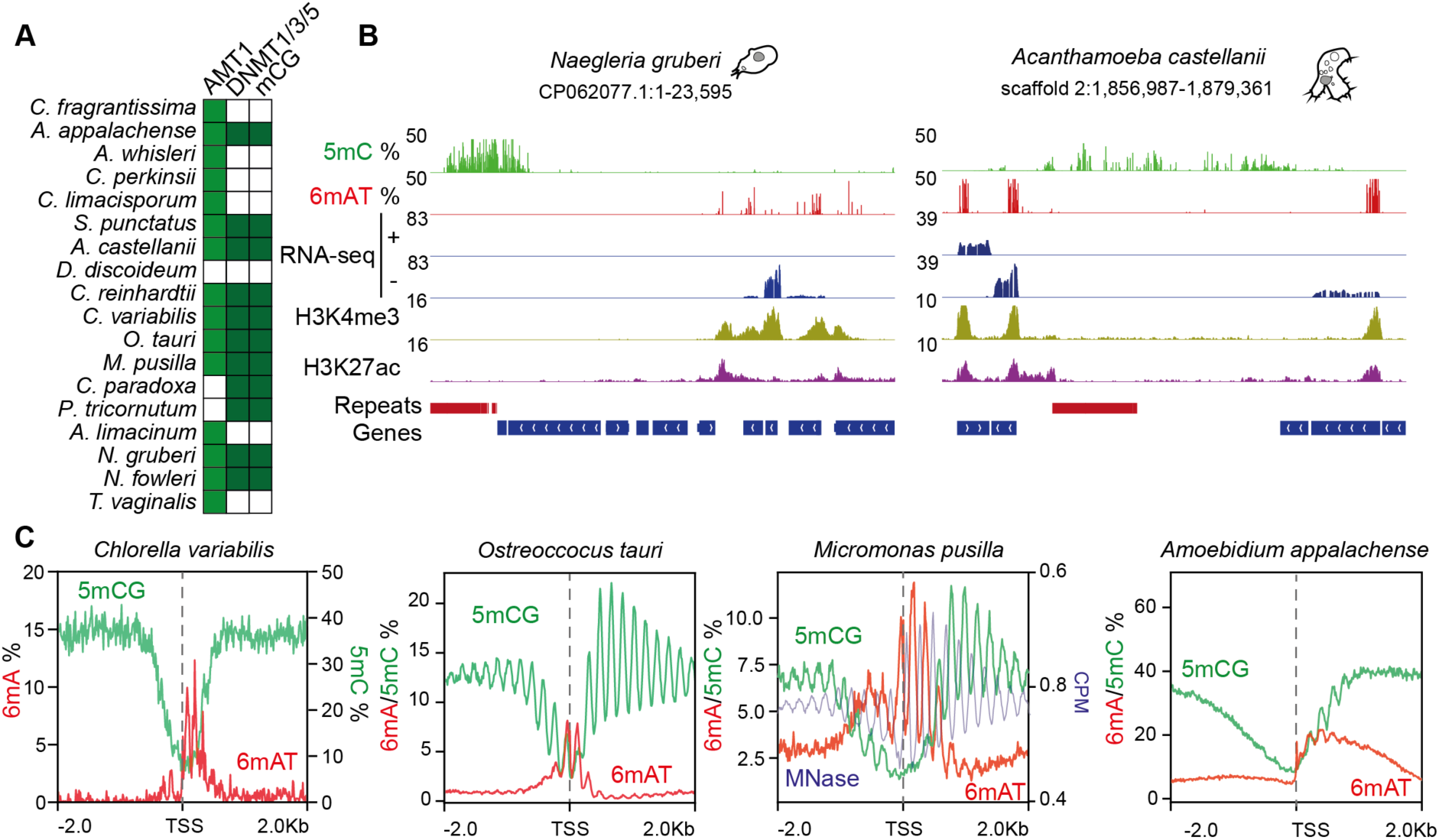
6mA and 5mC are compartmentalised in eukaryotic genomes. (**a**) Distribution of DNMTs and presence of 5mC in the CG context across the species in our dataset. Genome browser snapshots of (**b**) *Naegleria gruberi* and *Acanthamoeba castellanii* showing 6mA enrichment downstream of expressed genes and 5mC demarcating transcriptionally silent repetitive regions. RNA-seq (strand-specific) and ChIP-seq data is scaled using Counts per Million (CPM). (**c**) Average methylation levels centred at the TSS in species that have 5mC gene body methylation. For *Chlorella variabilis*, 6mA and 5mC axes are shown separately. For *Micromonas pusilla*, MNase data is shown to highlight the anti-correlation with both 5mC and 6mA peaks.

The recurrent loss of 5mC in eukaryotes is partly attributed to its mutagenic effects^42^. Species with elevated 5mC levels, like vertebrates, often have fewer CpG dinucleotides than expected by chance, since deamination of 5mC into thymines depletes CpGs over evolutionary time^43^. In contrast, animals with low or no 5mC lack this compositional bias^44^. Using a similar approach, we calculated observed versus expected compositional biases of ApT dinucleotides across our species panel, considering the AT context higher methylation. Global AT methylation levels did not correlate with ApT dinucleotide depletion, with a relatively constant ratio across species (**Extended Data Fig. 5**). Notably, CpG depletion was observed in species with very low 5mCG levels (e.g., *Naegleria*), while prasinophytes exhibited the opposite trend, with high 5mCG levels and enriched CpG composition (**Extended Data Fig. 5**), consistent with previous reports for this algal lineage^3^. These findings suggest that neither 5mC nor 6mA significantly shapes genomic base composition across the vast diversity of eukaryotes, in contrast with prior observations for 5mC in animals and plants.

One possibility is that the mutagenic effects and depletion of 6mA are predominantly observable in the genomic regions where this modification is enriched, specifically the post-TSS region of methylated genes. For instance, in invertebrates with 5mC on gene bodies, methylated genes tend to have lower CpG ratios than unmethylated genes^45,46^. To investigate this, we categorised the genes of each species based on the presence or absence of methylated ApTs and analysed the ApT dinucleotide density in those regions. In most species, there is no clear association between 6mA and local ApT composition (**Extended Data Fig. 6**). Surprisingly, in some species, methylated genes displayed higher ApT densities than unmethylated genes, contrary to what has been observed for 5mC methylation. Consequently, it appears unlikely that the recurrent loss of 6mA in eukaryotes is a consequence of its mutagenic potential, which seems to be limited.

Among the species in our dataset exhibiting both 6mA and 5mC, we observe two predominant patterns. A pattern in which 5mC is confined to transposable elements, while 6mA is enriched at expressed genes (**Fig. 3b**), suggesting potentially antagonistic functions in promoting or repressing transcription is found in *Acanthamoeba*, *Naegleria*, *Spizellomyces*, or *Chlamydomonas*. Nevertheless, in several eukaryotes like animals and plants, 5mC is also found within gene bodies^47,48^, but also the three chlorophytes and the ichthyosporean *Amoebidium appalachense* in our dataset^3,41^. In this second pattern found in these four species, 6mA predominantly occupies the five prime ends of the gene body, while 5mC peaks when 6mA diminishes (**Fig. 3c**). In all four species there is a distinct periodic 6mA pattern suggestive of internucleosomal linker positioning (**Fig. 3c**). To validate this observation, we used Micrococcal Nuclease sequencing (MNase-seq) data for *Micromonas pusilla*, confirming that 6mA peaks are anti-correlated with nucleosome positioning (**Fig. 3c**, **Extended Data Fig. 7**)^3^. In *Amoebidium*, both modifications coexist in the first three or four nucleosome territories downstream of the TSS(**Fig. 3c**, **Extended Data Fig. 7**). In the case of *Micromonas* and *Ostreococcus*, our Oxford Nanopore data aligns with the previously reported pattern of periodic 5mC internucleosomal distribution along the gene bodies of prasinophytes^3^. The original study already noted that periodic 5mC is weakly associated with transcription, suggesting a potential role in genomic compaction for these microalgae with some of the smallest eukaryotic genomes (e.g., *Ostreococcus tauri*, 12.6Mb)^3^. Importantly, we found that 6mA replaces 5mC in the first nucleosomes downstream of the TSS of actively transcribed genes. This implies that 6mA is likely important for keeping those genes transcriptionally active, as not expressed genes have less 6mA (**Extended Data Fig. 3**).

### Nucleosomes with H3K4me3 co-localise with 6mA

Our findings align with prior reports indicating an enrichment of 6mA in internucleosomal linker regions^15–17,31^. In ciliates, the deletion of AMT1 disrupts 6mA patterns, resulting in more diffuse nucleosome patterns around the TSS^17,27^. Most species in our dataset exhibit a periodic 6mA pattern, consistent with this suggested role. However, *Naegleria*, *Acanthamoeba*, or *Chromosphaera* lack a discernible periodic 6mA pattern (**Extended Data Fig. 6**). The absence of periodicity might be attributed to irregular nucleosome positioning in these species or potential issues with gene start annotations. However, we could not detect an apparent periodic pattern over individual genes, unlike the one observed in species displaying clear periodicity (**Fig. 3b**). Moreover, there is considerable variation in the average number of nucleosomes covered in 6mA territories across species. For instance, the stramenopile *Aurantiochytrium limacinum* displays only three narrow peaks, while the metamonad *T. vaginalis* exhibits constant 6mA periodic peaks across entire gene bodies, albeit with decreasing intensity (**Extended Data Fig. 6**). Thus, the connection between 6mA and nucleosomes is a consistent pattern across AMT1-encoding eukaryotes but presents subtle variations across lineages.

To gain further insights into what determines 6mA patterns and its association with nucleosome positioning, we conducted ChIP-seq for two histone modifications commonly linked to the TSS region and transcription: histone 3 lysine 4 trimethylation (H3K4me3) and lysine 27 acetylation (H3K27ac)^27,49^. We generated new data for four species and acquired publicly available datasets for two additional species^50^, ensuring representation across the most divergent eukaryotic groups. Both histone marks exhibited the anticipated patterns across eukaryotes, displaying enrichments post-TSS (**Fig. 4**)^27,51^. Subsequently, we categorised genes in each species based on the H3K4me3 signal. Remarkably, we found that 6mA mirrors the genomic regions coinciding with this histone mark (**Fig. 4**). Not only does the intensity of 6mA intensity align, but the width of deposition also largely corresponds across both epigenomic marks.

**Fig. 4.**
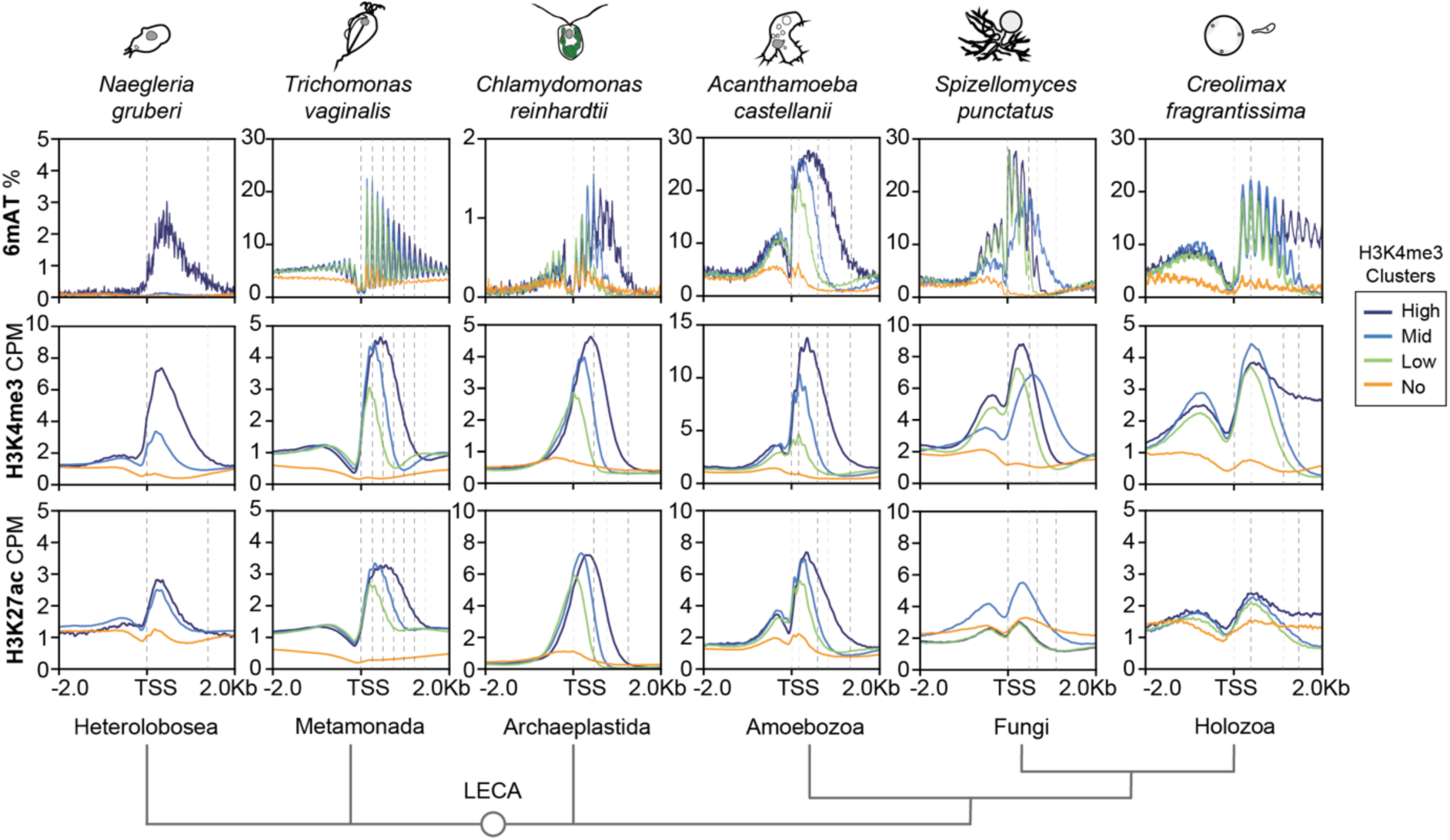
Nucleosomes with H3K4me3 modification recapitulate 6mA patterns. Average 6mA (on ApT context), H3K4me3 and H3K27ac across 5 eukaryotic lineages. For each species, genes have been clustered using the H3K4me3 signal, forming 4 clusters: high, middle, low and no signal. For *Naegleria gruberi*, since the amount of methylated genes is small, the classification was done independently, dark blue corresponds to 6mA methylated genes, pale blue corresponds to genes with H3K4me3 with no 6mA, and orange for genes that lack both H3K4me3 and 6mA. Dashed lines display the TSS, 6mA peaks (predicted internucleosomal linkers), and the boundaries of H3K4me3 signal across the 3 epigenomic marks.

In the ichthyosporean *C. fragrantissima*, we observed a potentially unique feature: some genomic regions were entirely covered by H3K4me3, spanning entire genes beyond the typical post-TSS restricted patterns (**Fig. 4, Extended Data Fig. 8**). Within these H3K4me3 domains, 6mA is consistently present throughout (**Extended Data Fig. 8**), strongly suggesting a tight link between the deposition of H3K4me3 and 6mA in these eukaryotes.

The other two species displaying slightly divergent patterns between 6mA and H3K4me3 are the heterolobosean *Naegleria gruberi* and the metamonad *T. vaginalis*. In *N. gruberi*, a few hundred genes show 6mA, so we categorised genes based on the presence or absence of 6mA and/or H3K4me3. Genes with 6mA, on average, showed the highest H3K4me3 signal (**Fig. 4**). However, a substantial number of genes displayed H3K4me3 without 6mA, suggesting that H3K4me3 is not dependent on 6mA (**Fig. 4**). This indicates that 6mA must have cues other than H3K4me3 for its deposition. In contrast, the H3K27ac signal was comparable between genes with 6mA and those with only H3K4me3.

*T. vaginalis* exhibits an opposite pattern. In this species, H3K4me3 and H3K27ac correlate with 6mA intensity, but 6mA extends beyond the initial modified nucleosomes (**Fig. 4, Extended Data Fig. 8**). This suggests that in *T. vaginalis*, both 6mA and H3K4me3 are dependent on transcriptional initiation. However, 6mA is deposited beyond H3K4me3, spanning broader regions. Overall, our data suggests that 6mA deposition linked to the AMT1 pathway is associated with nucleosome positioning across eukaryotes, particularly those linked with transcriptionally associated modifications. However, a potential hierarchy between these two chromatin components is likely species-specific, or perhaps is simplified in certain lineages.

## Discussion

We demonstrate the widespread presence of 6mA as a DNA base modification across eukaryotes, tracing its origin to the last eukaryotic common ancestor (LECA). Our taxon sampling encompasses previously unstudied eukaryotic diversity, and is consistent with the AMT1 pathway being present in early eukaryotes, irrespective of the debated root positions of the eukaryotic tree of life^52–54^. Unlike 5mC DNMTs acquired by LECA through independent bacterial transfers^55^, the MT-A70 family of 6mA methyltransferases was inherited in a single event from a bacterial donor (**Fig. 5a**). Pre-LECA, MT-A70 underwent duplications into six families, undergoing subfunctionalisation in heterodimeric partnerships and substrates, including RNA and DNA methylation. Post-diversification, MT-A70 families remained relatively static in eukaryotic evolution, contrasting with DNMT families undergoing lineage-specific duplications and expansions (Dim-2/RID in fungi, CMT and DRM in plants)^3,4,56^. Concurrently, 6mA patterns remained relatively constant across eukaryotes retaining the AMT1 pathway, consistently enriched in ApT dinucleotide context and associated with TSS. Conversely, 5mC displays evolutionary variability, ranging from gene body methylation to transposable element silencing or internucleosomal positioning, also methylating diverse sequence contexts beyond CpGs ^4–6^. As 5mC is presumed to ancestrally silence transposable elements^4,6,48^, its faster evolutionary dynamics may result from adaptations to track recurrent transposon invasions, contrasting with a slower concerted co-evolution of the 6mA-AMT1 pathway targeting endogenous genes. Our survey, with species from vastly divergent lineages showing very similar patterns, suggests LECA had a dual methylation system, with 6mA linked to transcription and 5mC likely associated with silencing (**Fig. 5a**).

**Fig. 5.**
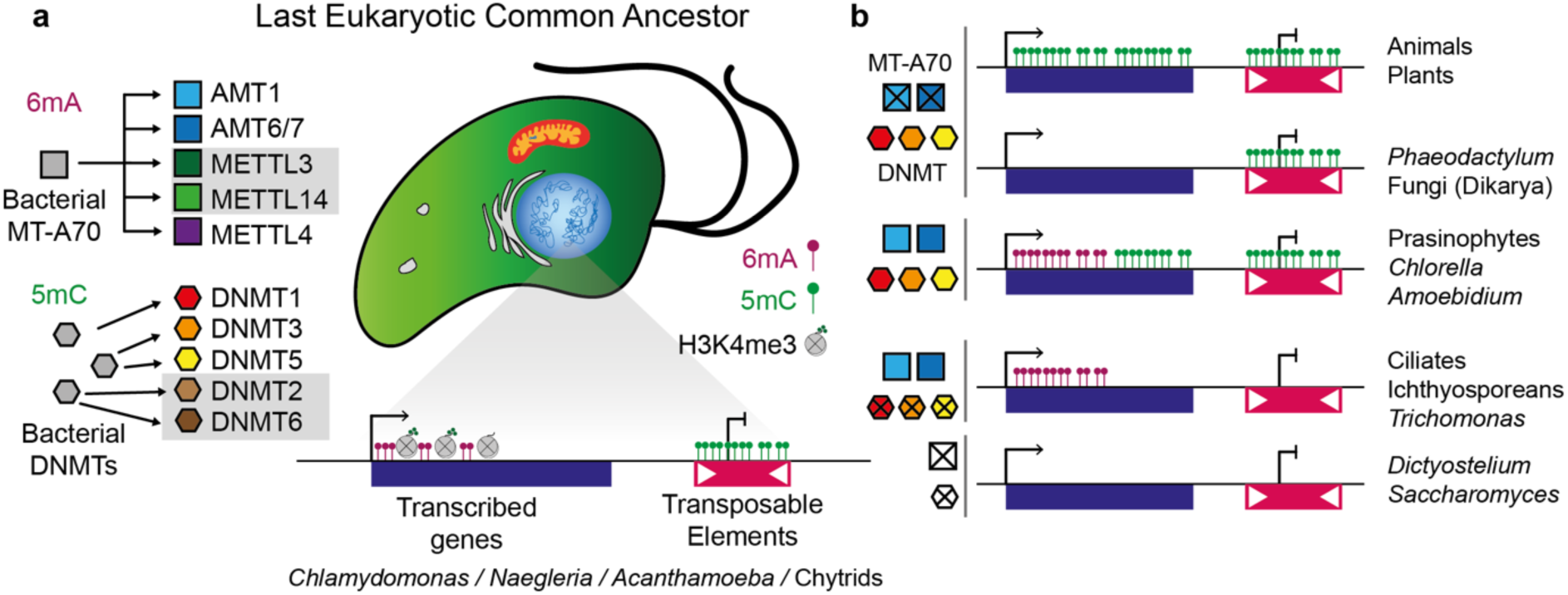
Reconstruction of the Last Eukaryotic Common Ancestor epigenome. (**a**) Diagram showcasing the distinct evolutionary origins of 6mA (MT-a70) and 5mC (DNMTs) enzymes in the advent of the LECA, and the inferred ancestral epigenome based on distribution and proposed roots of the eukaryotic taxa. Species genus represent examples of extant eukaryotes that still present the inferred ancestral patterns. (**b**) Derived epigenome patterns in modern eukaryotes associated with simplification in methyltransferase repertoires, with example species / lineages for each pattern. Crossed rectangles or hexagons indicate loss of the enzyme, which are colour / shape coded as in panel **a**. Presence of DNMTs does not imply co-occurrence of all 3 enzymes, but any of them.

Our study confirms Oxford Nanopore sequencing as a reliable platform for detecting genomic 6mA^32,57^, showcasing robust and reproducible patterns across species with diverse genome compositions. Long-read base pair resolution maps avoid pitfalls associated with other methods^23^, exemplified by the metamonad *T. vaginalis*. A previous study on *T. vaginalis*, reliant on antibody-based pull-downs, reported 6mA enrichment in transposable elements^58^. However, more than one third of the *T. vaginalis* genome derives from recently duplicated large Maverick DNA transposons^59^, making short read mappability particularly hard in this species, and repetitive DNA is prone to enrichment artefacts^24,60^. Instead, our *T. vaginalis* Oxford Nanopore data, reveals a strong periodic 6mAT pattern on transcribed genes without enrichment in repeats, aligning with patterns found in other eukaryotes in this study and others^15–17^. Nanopore technology could provide orthogonal confirmation to conflicting results from PacBio-based CLR methods, especially in cases like *Caenorhabditis elegans* where there is great debate on 6mA presence and potential function^10,19,61,62^. Moreover, in *C. elegans* the suggested 6mA methyltransferase, METL-9, belongs to a different enzyme family to MT-A70, with a distinct sequence motif (GGAG) instead of ApT^61,62^. It is possible that alternative 6mA DNA pathways may exist in eukaryotes, yet the few thousand 6mA sites detected in nematodes are not highly reproducible across samples and lack the strong clustered genomic patterns observed in AMT1-encoding species ^62^, but this could be the consequence of lacking a maintenance type methyltransferase. The nucleotide salvage pathways incorporating 6mA from RNA and mitochondrial DNA into nuclear DNA, or RNA:DNA hybrids might explain some of these off-target effects in previous reports^21–23,63^. Conversely, the robust and reproducible detection of symmetrical ApT methylation in AMT1 encoding eukaryotes implies a functionally significant and stable component of chromatin.

Our discovery about the recurrent loss of the AMT1 6mA pathway in eukaryotes challenges previous assumptions in DNA methylation evolution. Similarly to DNMTs and the RNA 6mA METTL3/14 pathway, the 6mA DNA pathway is frequently lost across eukaryotes ^4,41^. However, 6mA does not have an obvious mutagenic effect; so this pattern may suggest that 6mA loss reflects evolutionary contingency, indicating that its function can be compensated through alternative means without severe consequences. These losses have resulted in a wide diversity of patterns and combinations of 6mA and 5mC in extant eukaryotes (**Fig. 5b**).

The function of 6mA in nucleosome positioning remains unclear, as ciliates with mutated AMT1 exhibit disordered nucleosomes yet remain generally viable^17,27^, primarily only lethally affected in sexual reproduction. In ciliates, the effect on nucleosome positioning is also stronger *in vitro* than *in vivo*^17,27,31^. Notably, some of our surveyed species lack a periodic 6mA pattern, casting doubt on the importance of 6mA in preventing nucleosome movement as a universal feature. However, our data does suggest an ancient link between 6mA and H3K4me3 demarcating the 5 prime end of genes, possibly together with the histone variant H2A.Z^27^. Still, considering the prevalence of H3K4me3 as a post-TSS nucleosome marker in eukaryotes^49^, including those lacking 6mA, this implies that 6mA may not be essential or is easily substituted or compensated in this genic region. Of note, H3K4me3 and H2A.Z usually anti-correlate with the presence of 5mC in a vast diversity of eukaryotes^48,64,65^, thus the link between H3K4me3 with 6mA might help in this chromatin compartmentalisation. Moreover, in thinking about the origin of eukaryotic nucleosomes^66^, 6mA being found in the early internucleosomal linkers should be taken into account. Biophysically, 6mA is believed to decrease double-strand DNA stability, an effect potentially advantageous for transcription at the start of highly expressed genes^21,26^. However, to understand the 6mA role as an epigenetic mark in eukaryotes, the identification of potential readers is necessary. Drawing parallels with 5mC, it is known that 5mC increases DNA stiffness^21^, yet the role of 5mC across eukaryotes likely depends more on the “readers” of the base modification than its intrinsic biophysical properties^67^.

Intriguingly, 5mC is predominantly retained in major multicellular eukaryotic lineages (plants, animals, and Dikarya fungi), whereas the AMT1 6mA pathway was lost in these lineages. In plants and animals, 5mC evolved to methylate the entire gene body (**Fig. 5b**)^4,5,48^, replacing 6mA in the post-TSS region, yet this substitution is unlikely the sole cause for 6mA loss in these lineages. Attesting this, we show examples of unicellular relatives of plants and animals with 5mC gene body methylation coexisting with 6mA, indicating that probably 6mA and 5mC play distinct roles even if found in the same regions (**Fig. 5b**). In conclusion, our data reshapes the understanding of 6mA function and evolution in eukaryotes, emphasising that the two forms of DNA methylation are integral components of the original chromatin toolkit of eukaryotes. Although ciliates have spearheaded our understanding of 6mA in eukaryotes, their drastically unusual genome organisation, with a micronucleus and a macronucleus, and a lineage-specific expansion of MT-A70s^17,27^, will benefit from complementary insights from alternative lineages. This expanded taxon sampling in potentially tractable systems with canonical eukaryotic genomes will drive new efforts to uncover 6mA functions and elucidate why this crucial pathway was lost in major multicellular lineages.

## Materials and Methods

### Sequence search and phylogenetic analysis

A collection of eukaryotic proteomes spanning the broadest diversity of lineages was scanned using HMMER3 with the PFAM domain for MT-A70^68^, using an e-value of 0.0001 as a threshold. Additionally, AMT6/7 sequence from *Tetrahymena termophila* was used as a query against the same database using BLASTP, to obtain divergent orthologues that were filtered out with the hmmsearch approach. In parallel, *Tetrahymena termophila* MT-A70 AMT1 and AMT7 sequences were used to BLASTP against NCBI “nr” databases excluding eukaryotes as taxonomic hits. The resulting proteins were merged into a multisequence alignment using MAFFT L-INS-i mode^69^. The alignment was trimmed with trimAl -gappyout parameter^70^, and a maximum likelihood phylogeny was obtained using IQ-TREE^71^, with automatic model selection, computing 1000 ultrafast bootstrap and ALRT replicates as nodal supports. To determine the distribution of MT-A70 in prokaryotes, the same BLASTP approach was used allowing 250 hits against NCBI “refseq_select” database, specifying bacteria and archaea (separately). To determine the presence of p1 and p2 orthologues, we used BLASTP against the custom eukaryotic proteome database, requiring an e-value < 0.0001.

### Culture of protists and genomic DNA extraction

*Creolimax fragrantissima, Abeoforma whisleri* and *Corallochytrium limacisporum* cells were grown axenically in marine broth liquid medium (Difco 2216) at 17°C, 17°C, and 23°C respectively. *Chromosphaera perkinsii* cells were grown axenically at 23°C in liquid medium (containing 3g of yeast extract, 3 g of malt extract, 5 g of peptone, 10 g of glucose and 20 g on NaCl per litre of distilled water). *Acanthamoeba castellaniii* cells were grown axenically at 23°C in ATCC medium 712. *Naegleria gruberi* cells were grown axenically at 30°C in ATCC medium 1034. *Aurantiochytrium limacinum* cells were grown axenically at 19°C in ATCC medium 790. *Spizellomyces punctatus* cells were grown axenically at 17°C in liquid medium (containing 2.5 g of yeast extract, 0.5 g of K_2_HPO_4_, 2.5 ml of ethanol, 15 ml of glycerol and 485 ml of Milli-Q water). *Dictyostelium discoideum* was grown axenically at 23°C in liquid HL5-C medium (Formedium).

We performed DNA extractions for *N.gruberi*, *D.discoideum*, *C.perkinsii, A. whisleri* and *S.punctatus* from confluent cultures grown into 25 ml flasks for 7 days, and 4 days for *C.limacisporum*. For *A.limacinum* cells grown for 7 days and then 1 ml was passed into two 25 ml flasks with fresh medium to enrich for zoospores and then collected two days later. In all cases, cells were centrifuged at 4400 rpm for 5 min and the supernatant discarded before the DNA extraction.

For *A. castellani* trophozoite DNA, we obtained cells from a confluent 7 day culture. For the cystic stage, cells were grown for 5 days in 25 ml flasks and then media was removed and replaced by 5 ml of encystment medium (containing 3.728 g of KCl, 1.68 g NaHCO_3_, 0.986 g of MgSO_4_ x 7H_2_O, 0.03 g of CaCl_2_ x 2H_2_0 and 0.017 g of 2-Amino-2-methyl-1,3- propanediol per 500 ml of distilled water)^72^, after 3 days of incubation at 23 °C *A.castellani* cysts were collected for DNA extraction.

For *C.fragrantissima* cells were grown until confluency for 5 days in 25ml flask, then cells were scratched and passed into 50 ml flask with 25 ml of fresh medium. These new flasks were grown for 48h under gentle agitation at 17°C and then were filtered using a 20μm cell strainer (pluriSelect) and collected into a 50 ml Falcon tube to separate the amoebas from the mature coenocytes, as described previously^39^. Both amoebas and coenocytes were collected for independent DNA extraction.

To break the cell wall of *S. punctatus* and *C. fragrantisiima* heat shock was applied, the cells were frozen with liquid nitrogen and then thawed at 60°C, repeating this 3 times. For *C. perkinsii* and *A. castellani* cysts we immersed the pellets in liquid nitrogen immersion followed by grinding with pestle and mortar.

The resulting cell pellets for *N.gruberi, A.castellanii* amoebas and cysts*, D.discoideum*, *C. perkinsii*, *C. limacisporum*, and *A. limacinum* were used for DNA extraction using the Qiagen MagAttract HMW DNA kit following the manufacturers whole blood protocol. For *S. punctatus* and *C. fragrantissima* NEB Monarch Genomic DNA Purification Kit was used following the animal tissue protocol. For *A. whisleri* we used a phenol/chloroform genomic DNA extraction method.

The genomic DNA of *T. vaginalis* strain G3 was obtained from ATCC, and the DNA of *M. pusilla* CCAP 1965/4*, C. variabilis* NC64A and *C. paradoxa* CCAP 981/1 were obtained from CCAP (Oban Culture Collection of Algae and Protozoa).

### Nanopore sequencing, basecalling and methylation analysis

We quantified genomic DNA using a Qubit 3 Fluorometer with the dsDNA BR Assay Kit and assessed DNA size with a TapeStation 2200 using the Genomic DNA ScreenTape Assay. We then started with 1-1.5 μg of high molecular weight genomic DNA that was ligated with Nanopore SQK-LSK110 ligation kit following manufacturer’s instructions and sequenced later in MinION R9 flowcells (see details in **Supplementary Table 1**).

The resulting fast5 files were then used as input for Guppy v6.5.7, and the Rerio modified basecalling model “res_dna_r941_min_modbases-all-context_v001” was specified, aligning the reads to the reference genome (--align_ref) parameter. The list of reference genomes is found in **Supplementary Table 2**. The resulting BAM files were then merged and sorted using samtools, and then the Oxford Nanopore “modbam2bed” (0.9.1) was used to obtain either 6mA (-m 6mA) or 5mC (-m 5mC) basecalls. The resulting bedMethyl files were then used to extract the neighbouring bases for each A position, gathering 1 base upstream and 2 downstream using BEDTools^73^, and the stranded (5’ to 3’) four-nucleotide context was included for each position. Dinucleotide contexts were divided into AT, AA, AC and AG, and bigwig files were generated using UCSC bedGraphToBigWig tool. Visualisation of average methylation levels on genes, TSS and transposable elements was obtained using DeepTools2^74^.

Global methylation levels were computed in R, and the regional average levels of 6mA were computed using the Bioconductor bsseq package^75^, treating the ApT dinucleotides as if they were CpGs.

### RNA-seq analysis

Publicly available RNA-seq datasets for all species were downloaded from ENA from various studies (**Supplementary Table 3**). The data was mapped to the reference genomes using HISAT2^76^, restricting intron size to 10,000 bp. Stringtie was then used to obtain the Transcripts per Million measure for all gene models for each species, specifying the stranded information if the original libraries had strand-specific information ^77^.

### ChIP-seq library preparation and analysis

ChIP-seq was performed as previously described^78^ with modifications. Briefly, 100 ng chromatin per species was pooled per ChIP. Pooled chromatin was incubated at 4°C for 12−14 h with rotation with 2.5 µg anti-H3K27ac (Abcam, ab4729) or 2.5 µg anti-H3K4me3 (Millipore, 07-473). Immunoprecipitated complexes were captured using a mix of Protein A (16-661, Sigma-Aldrich) and Protein G magnetic beads (16-662, Sigma-Aldrich), washed, and reverse crosslinked for 30 min at 55°C followed by an hour at 68°C. Immunoprecipitated DNA was purified using SPRI beads (A63881, Beckman Coulter). The NEBNext Ultra II DNA Library Prep Kit (New England BioLabs) was used to produce the resulting ChIP libraries according to the manufacturer’s protocol.

The data for *C. reinhardtii* and *T. vaginalis* was downloaded from ENA, belonging to previous studies^50,79^, (see SRR numbers in **Supplementary Table 4**).

All ChIP-seq data was analysed using fastp to trim the reads, and these were mapped to the genomes using bowtie2, allowing a maximum insert size of 2000 base pairs (-I 2000). Duplicate reads were removed using Sambamba. DeepTools2 was used to generate bigwig files and to visualise epigenomic data, as well as Integrative Genome Viewer.

### Genome assembly and re-annotation

The genomes of *N. gruberi* and *C. perkinsii* were re-assembled for this study. We used the Oxford Nanopore reads and did conventional base calling using Guppy with the “sup” model and splitting chimeric reads. The resulting reads were then used as input for Flye (v2.9-b1768) using –nanopore_hq option, and allowing two steps of polishing^80^. Publicly available Illumina reads^81^ were used for polishing using HyPo^82^. For *N. gruberi*, the resulting contigs were scaffolded using *Naegleria lovaniensis* chromosome scale genome^83^ with RagTag^84^. Publicly available RNA-seq for each species^81,85^ was aligned to the genomes using HISAT2 with the -dta parameter with a maximum intron size of 10 kb, and Stringtie was then used to predict transcript models^76,77^. In parallel, Trinity was used to build de novo transcriptome assemblies^86^, which were mapped to the genome using gmap. The best transcripts were selected using Mikado^87^, which were then used to generate exon hints for Augustus^88^. Liftoff^89^ was used to move the annotations of the previous genome assemblies to the new assemblies^81,90^, and those were used as coding sequence hints. Then Augustus was trained on the best Mikado gene models for each species, and those species models were used to predict the genes with both exon and coding sequence hints. In the last step, transcript UTRs were added to the Augustus gene models using PASA with the best Mikado transcripts as input^91^.

For *C. limacisporum*, *C. variabilis*, *C. fragrantissima* and *A. whisleri*, the annotations were also updated with available RNA-seq data to include UTRs and improve TSS annotation^39,92,93^. We used a combination of Trinity and Stringtie as described above to generate a non-redundant transcript set with Mikado, which was then used to update gene structures with PASA.

Repeat annotations were generated for those species that lacked them, for which we used RepeatModeler2^94^ including the LTR finding module, and mapping the consensus repeat library against the genome using RepeatMasker.

## Acknowledgements

We would like to thank Iñaki Ruiz-Trillo and Meritxell Antó for sharing some of the initial cultures used in this study, and Batool Mahmoud for helping with cultures and DNA extractions. We would like to thank Omaya Dudin and Ozren Bogdanovic for critically reading the manuscript. This work used computing resources from Queen Mary University of London’s Apocrita HPC facilities. This work was funded by the Horizon 2020 Framework Programme to AdM (European Research Council Starting Grant action number 950230), PCR was funded by a QMUL PhD fellowship, CN was funded by FPI PhD fellowship from the Spanish Ministry of Science and Innovation, research in ASP group was supported by the European Research Council (ERC-StG 851647) and the Spanish Ministry of Science and Innovation grant (PID2021-124757NB-I00) as well the Spanish Ministry’s support to the EMBL partnership, the Centro de Excelencia Severo Ochoa and the CERCA Programme (Generalitat de Catalunya). V.S and E.C are supported by a grant (PID2020-120609GB-I00) by MCIN/AEI/10.13039/501100011033 and by a grant 2021SGR00751 of the Departament de Recerca i Universitats de la Generalitat de Catalunya.

## Contributions

AdM conceived and designed the study. PRC obtained nanopore sequencing for all but two species. PRC, VO, LS and AdM performed bioinformatic analysis. CN, DLA and ASP performed the ChIP-seq experiments. VS, EC provided nanopore data for *Abeoforma whisleri* and shared cultures. AdM and PRC drafted the manuscript, and all authors critically read and commented on the manuscript.

