## SupplementaryFigures for "Adenine DNA methylation associated to transcription is widespread across eukaryotes"

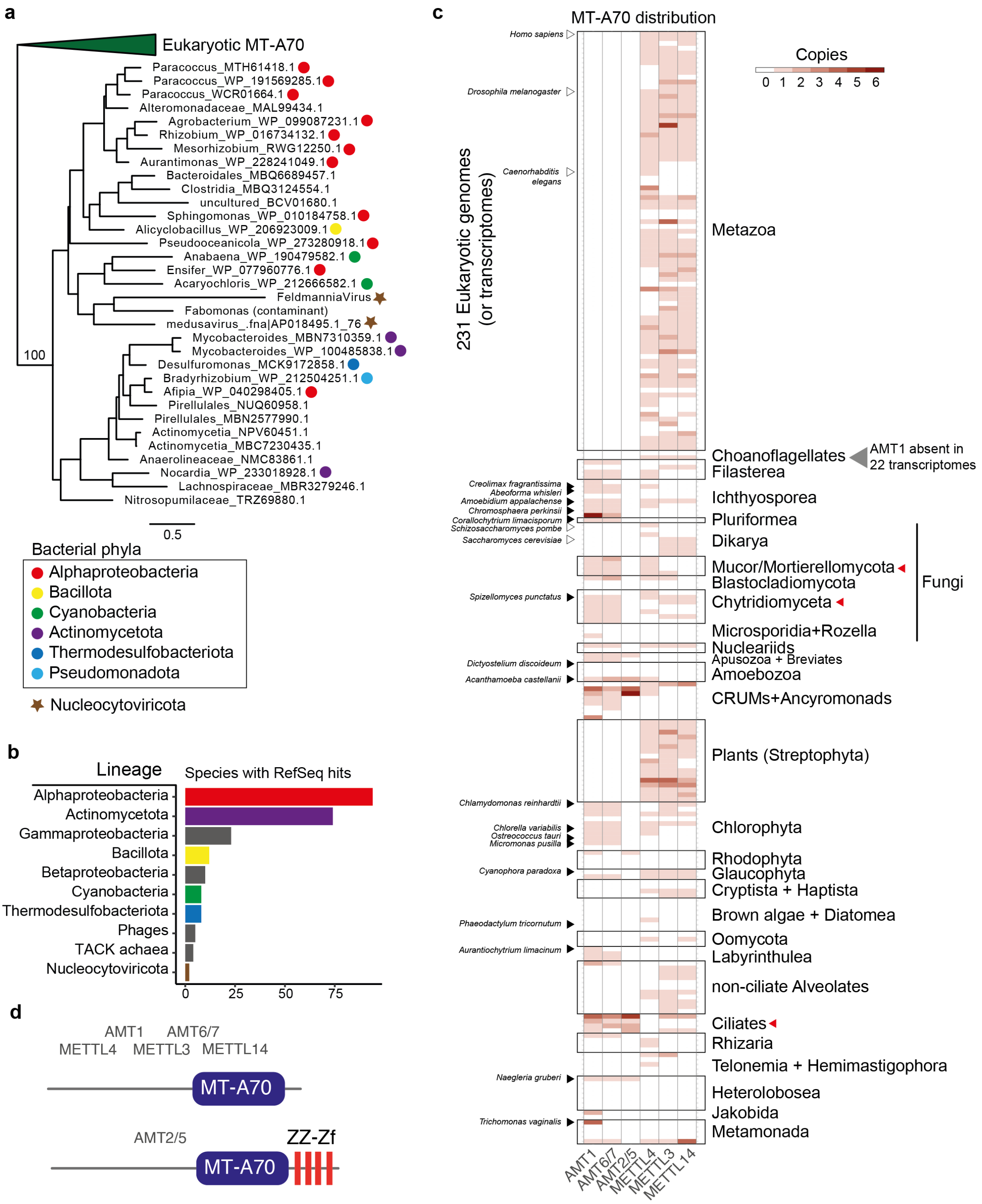


**Extended Data Fig. 1. Bacterial origins, structure and distribution of MT-A70**. (**a**) Detail of the maximum likelihood represented in Fig. 1B highlighting the prokaryotic outgroup, colour coded by bacterial lineage (for those bacteria with genus / species defined). (**b**) Taxonomy count of MT-A70 BlastP searches against NCBI RefSeq_select database - January 2024). (**c**) Heatmap showing the number of MT-A70 family members across a collection of 231 eukaryotic species covering a wide scope of known diversity. Classification obtained from a phylogenetic tree in **Fig. 1b**. Black triangles are the species sequenced in this study, and red arrows indicate lineages / species for which 6mA has been previously confirmed at robust high levels, white triangles indicate species for which 6mA is contested. (**d**) Domain architectures of eukaryotic MT-A70 families, defined with Pfam domains MT-A70 (PF05063) and ZZ (PF00569).

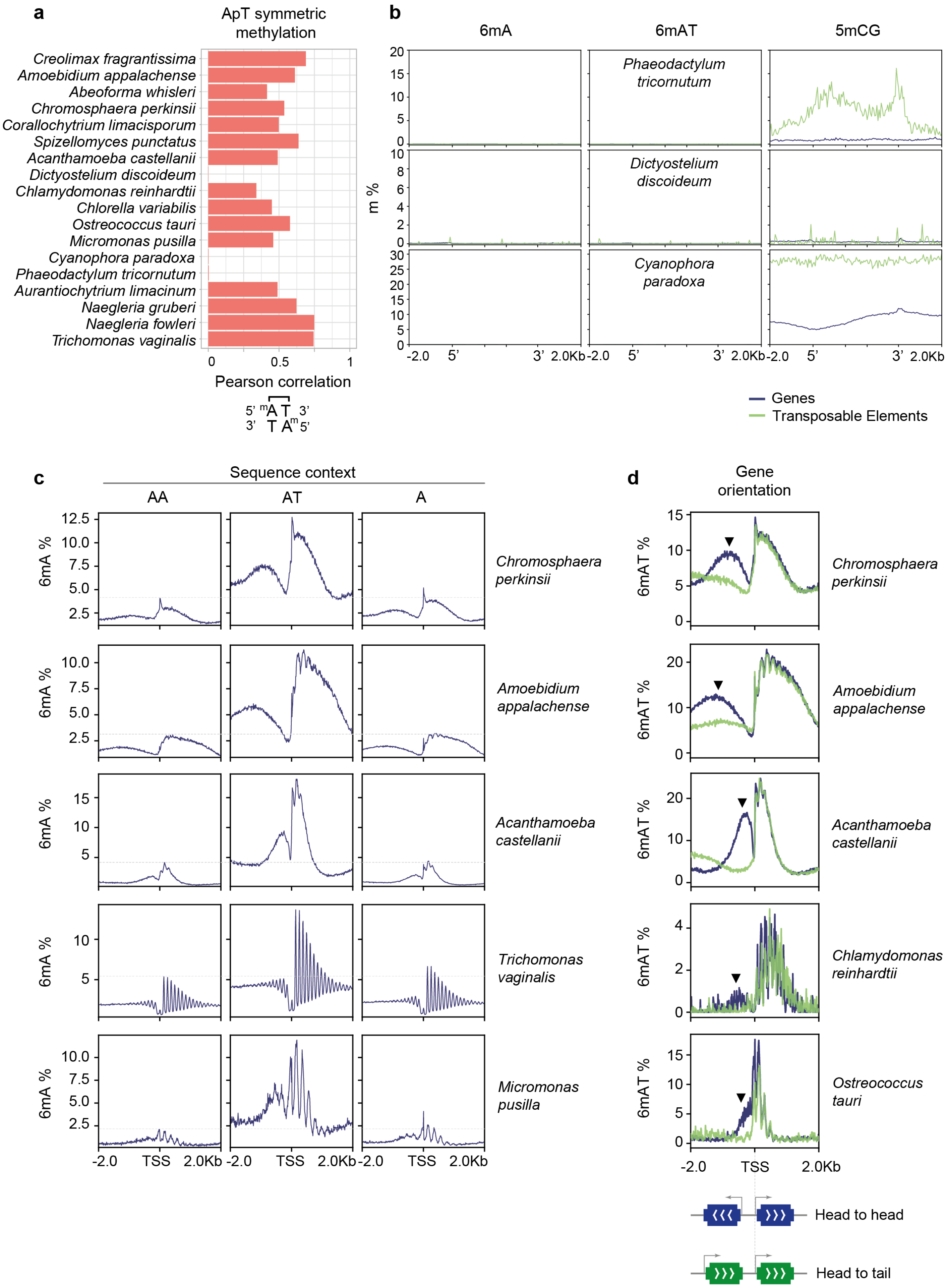


**Extended Data Fig. 2. Patterns of 6mA distribution in eukaryotes**. (**a**) Global correlation of 6mA levels in ApT dinucleotides. Pearson correlation was obtained for ApT positions for which there was at least 10x coverage in each strand. Note lack of correlation in AMT1-lacking species. (**b**) Average methylation levels in 6mA and 5mC for three species that lack AMT1 or AMT6/7. *Dictyostelium discoideum* also lacks DNMTs and has been previously reported to lack 5mC, thus the signal is likely noise (also driven by the low number of annotated TEs - 88). Borders of genes and TEs are represented by the 5' and 3' sites. The first 1500 bp post TSS and before the TES are unscaled. (**c**) Average 6mA levels around the Transcriptional Start Site (TSS) in five species with high 6mA levels (Fig. 2A), divided by sequence context. AT dinucleotides are the preferentially methylated context, but the AA context also gets residual methylation on the same regions, yet on average less than the total 6mA methylation. The dashed line highlights the max reached by AA methylation. (**d**) Average 6mA level plot around the TSS in species with marked "promoter" methylation patterns. Genes with a head to head orientation are displayed in blue and genes with a head to tail orientation are displayed in green. The black arrow shows the artifactual "promoter" peak driven by head to head oriented genes.

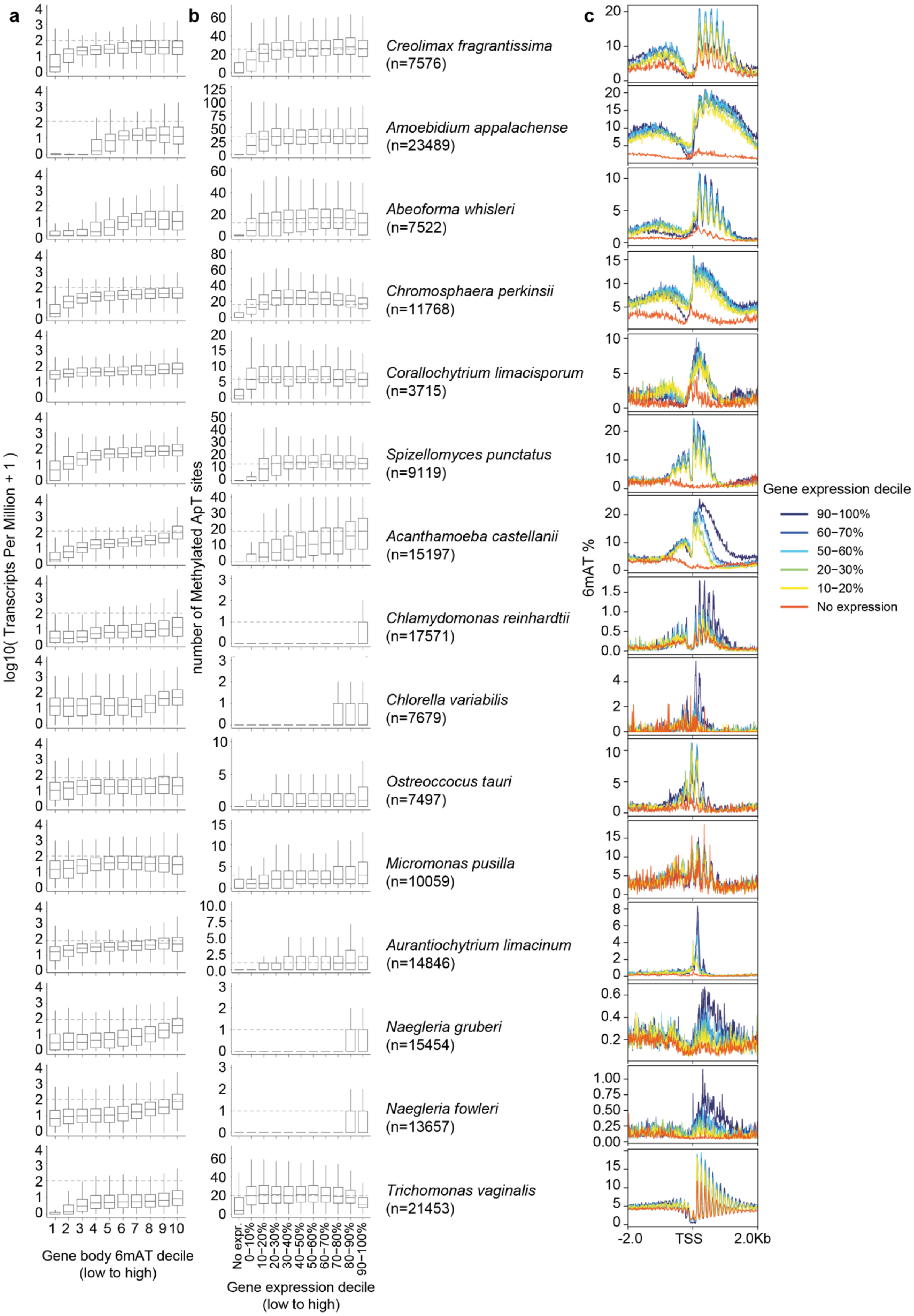


**Extended Data Fig. 3. 6mA correlates with higher transcription across eukaryotes encoding AMT1**. (**a**) Distribution of transcription levels of genes divided in 10 deciles sorted by the gene body 6mA levels in the AT context. (**b**) Distribution of methylated AT sites (>10%

6mA) in gene bodies sorted in 10 deciles by transcriptional level, where not expressed genes (TPM <1) are all shown separately. “n=” in parenthesis indicates the amount of genes shown for each species, which are selected for a minimal coverage (>4x). Centre lines in boxplots are the median, box is the interquartile range (IQR), and whiskers are the first or third quartile ± 1.5× IQR. (**c**) Average 6mA levels in the AT context around the TSS for genes sorted by transcriptional levels as in panel **b**. Decile of expression colour code shown in legend.

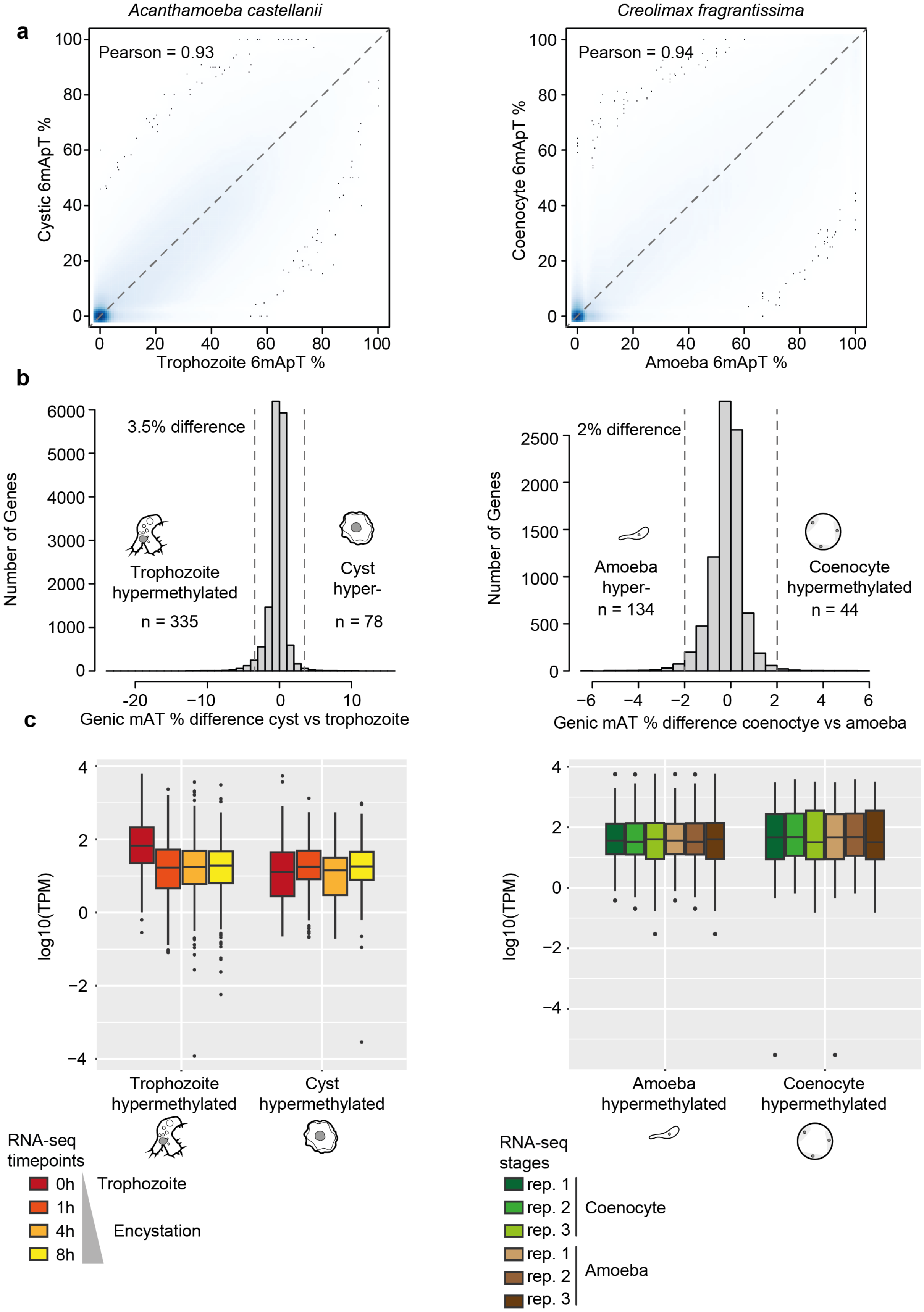


**Extended Data Fig. 4. 6mA methylation is modestly responsive to transcriptional changes**. (**a**) Comparison of 6mA levels in ApT dinucleotides across two stages of *Acanthamoeba castellanii* and *Creolimax fragrantissima*. Only ApT sites with at least 10x coverage in both samples are shown. (**b**) Distribution of genic 6mA methylation difference across stages. Dashed lines indicate the threshold of difference used to define hypermethylated genes in a given stage. (**c**) Boxplot displaying the transcriptional levels of genes with the variable 6mA levels from panel **b** across cell stages. Each RNA-seq sample is colour coded according to the legend, with datasets coming from previous publications[38,39]. Centre lines in boxplots are the median, box is the interquartile range (IQR), and whiskers are the first or third quartile ± 1.5× IQR.

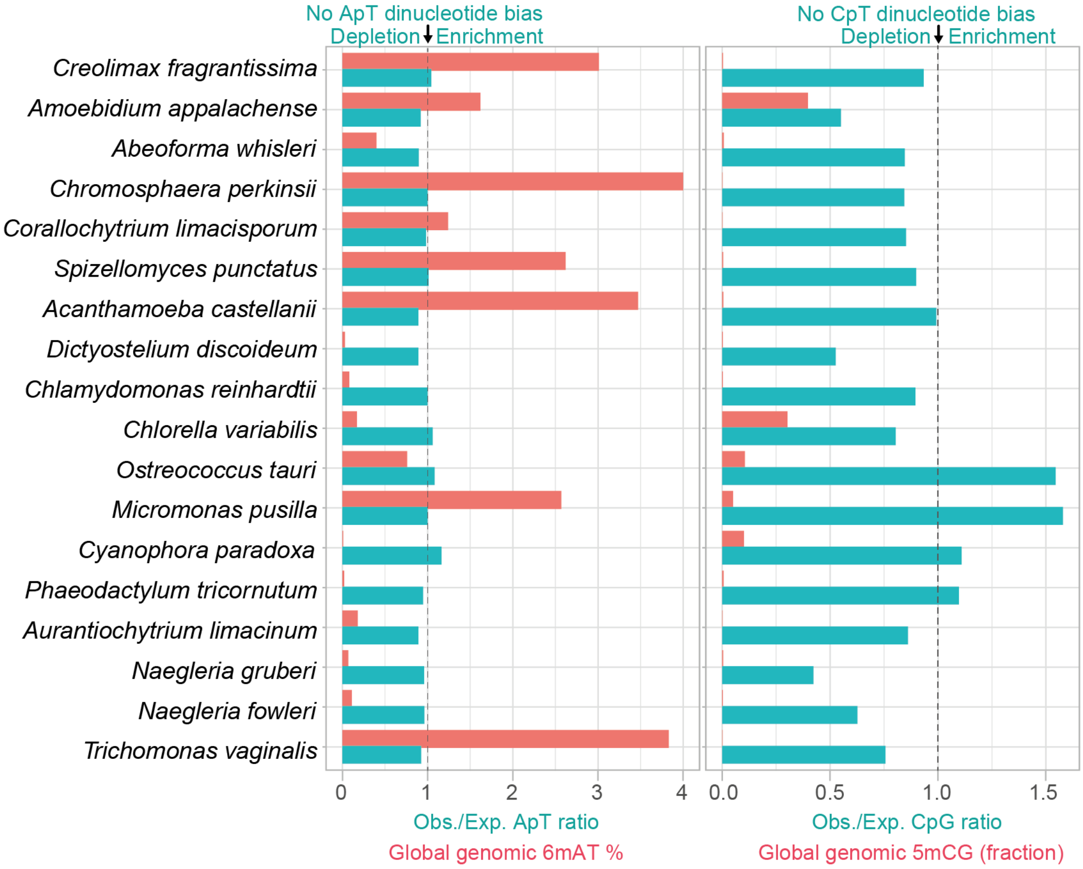


**Extended Data Fig. 5. 6mA methylation is not associated with a depletion of AT dinucleotides**. The global genomic 6mA levels in AT context (red) are shown next to the Observed versus Expected ApT dinucleotide ratios (turquoise) in each of sampled genomes. An Observed vs Expected ratio of 1 indicates ApT dinucleotides are found in the expected ratio given a genomic AT%, <1 levels show a depletion, whereas >1 levels indicate an enrichment. For comparison, global genomic 5mC levels (shown as a fraction with maximum being 1.0 in red) in the CG context are shown next to the Observed vs Expressed CG dinucleotide context ratio (turquoise) for the same set of species.

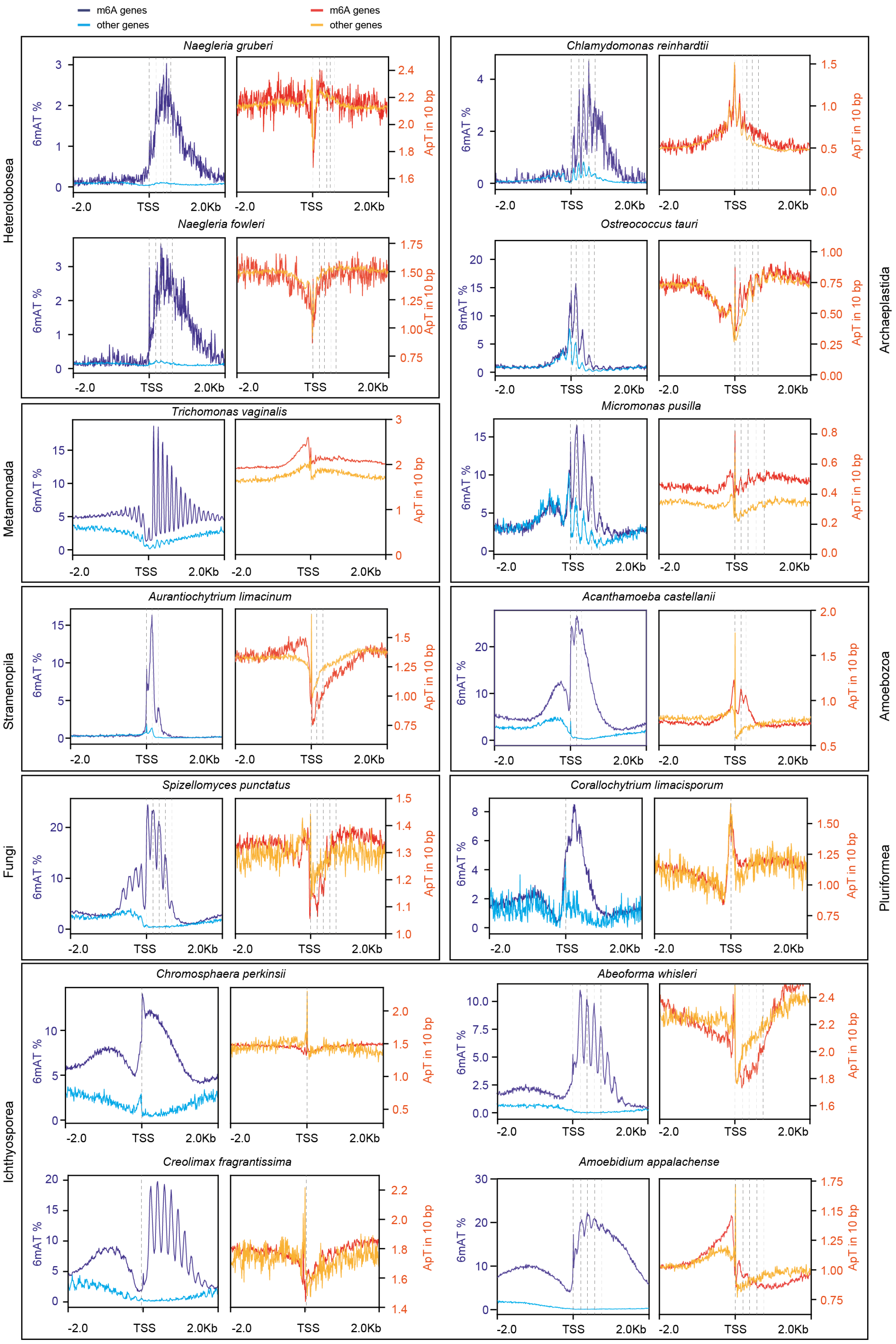

m6A genes
other genes
m6A genes
other genes
Heterolobosea
Naegleria gruberi
Naegleria fowleri
Metamonada
Trichomonas vaginalis
Stramenopila
Aurantiochytrium limacinum
Fungi
Spizellomyces punctatus
Ichthyosporea
Chromosphaera perkinsii
Creolimax fragrantissima
Archaeplastida
Chlamydomonas reinhardtii
Ostreococcus tauri
Micromonas pusilla
Amoebozoa
Acanthamoeba castellanii
Pluriformea
Corallochytrium limacisporum
Abeoforma whisleri
Amoebidium appalachense
6mAT %
ApT in 10 bp
-2.0
TSS
2.0Kb

**Extended Data Fig. 6. 6mA rarely recapitulates the ApT dinucleotide composition downstream of the TSS**. Average 6mA levels in the AT context shown side by side to average ApT content in genes separated by their methylation status. Methylated genes are shown in dark blue (6mA plot) and dark red (ApT content plot), whereas unmethylated genes are shown in pale blue (6mA plot) and orange (ApT content plot). Windows for averaging are 10 bp long. Enrichment of ApT in the TSS is likely driven by lack or inaccurate UTR annotations in many gene models, with 5’ end of genes being the start codon “ATG”.

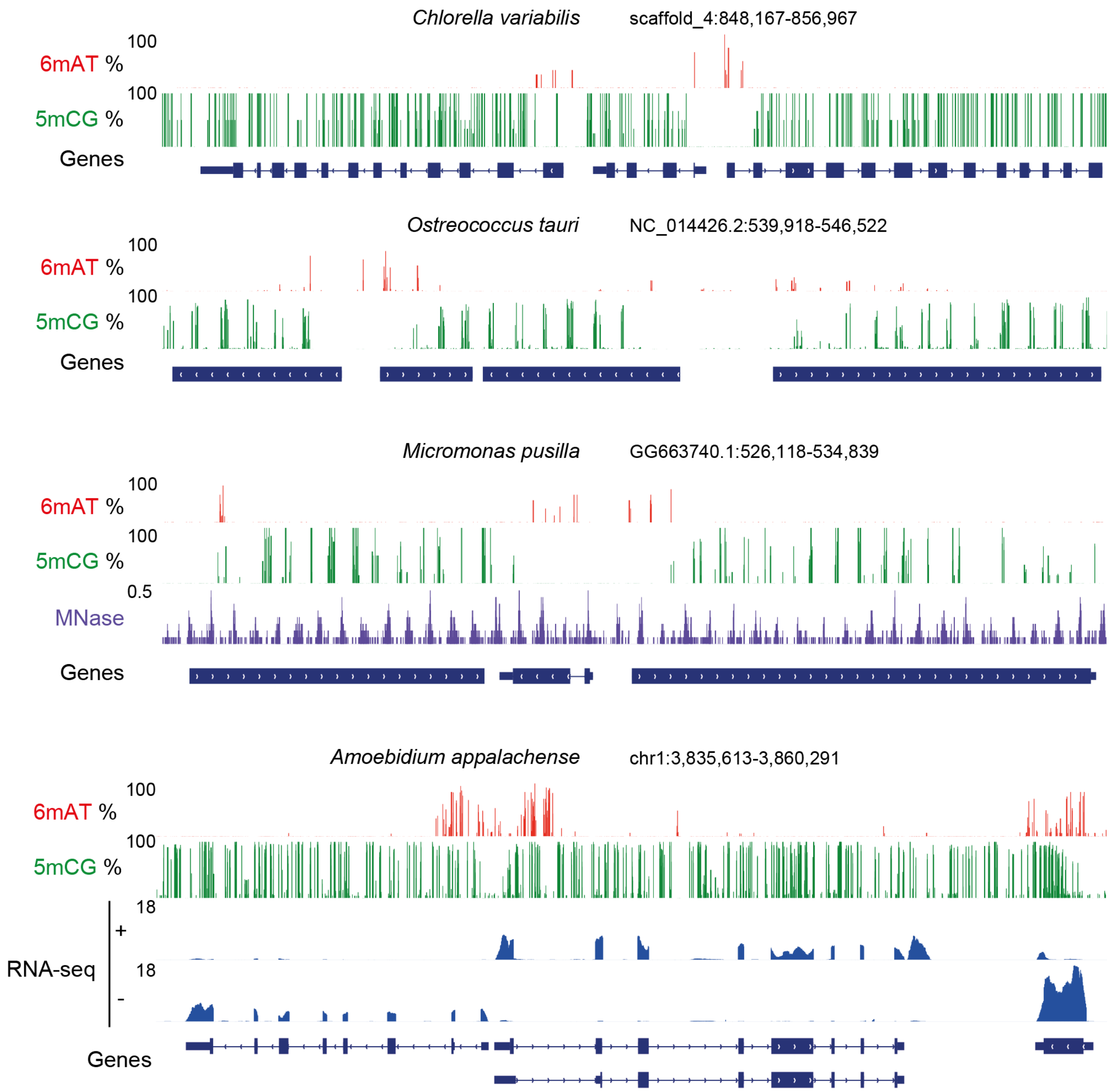


**Extended Data Fig. 7. 5mC in gene bodies coexists with post-TSS 6mA**. Genome browser snapshots of four species with 5mCG on gene bodies. MNase and RNA-seq data is normalised using Counts Per Million (CPM).

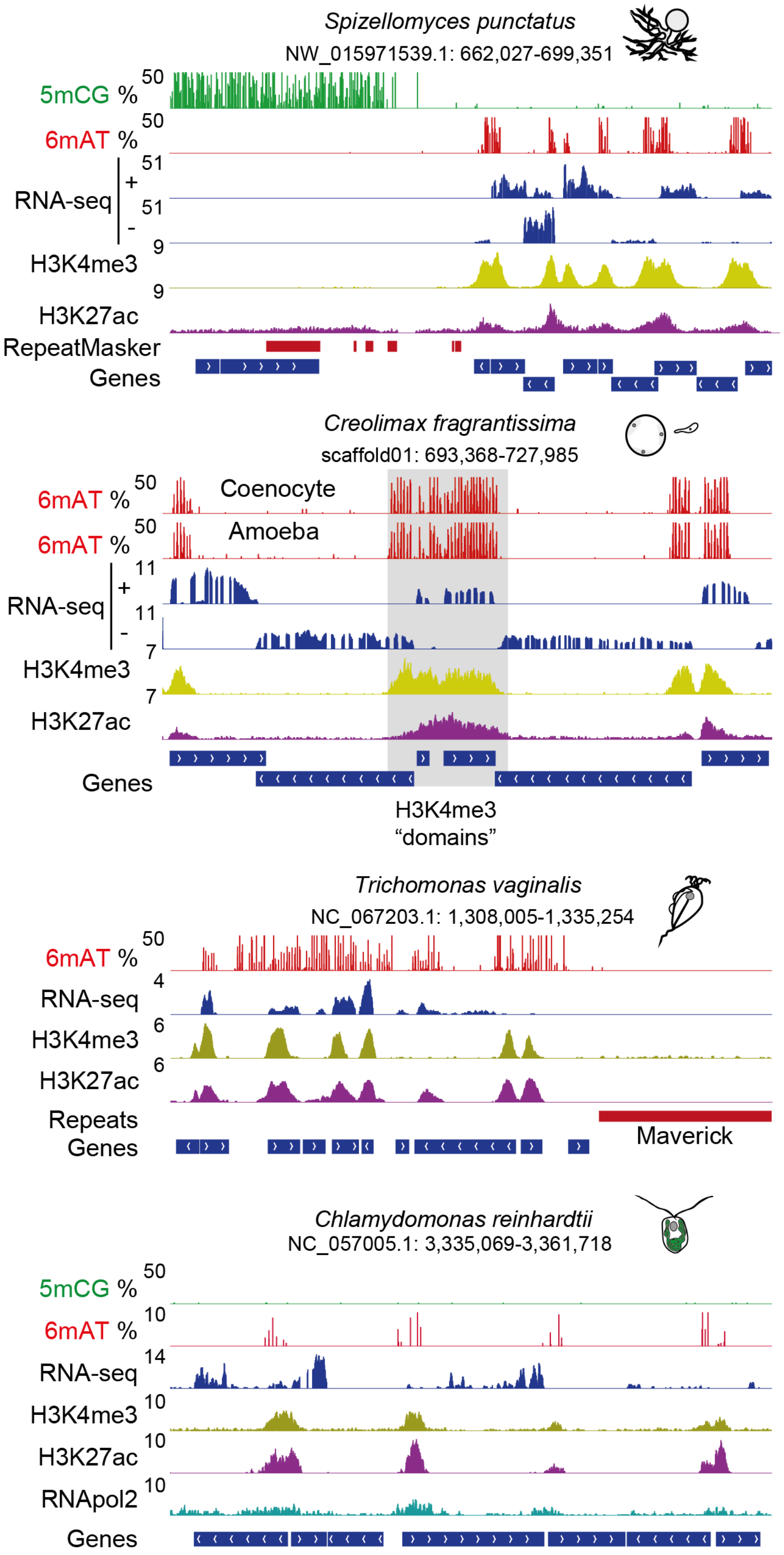


**Extended Data Fig. 8. 6mA is located in H3K4me3 demarcated regions**. Genome browser snapshots of four species with ChIP-seq data, with genomic coordinates on top. RNA-seq (positive and reverse strand) and ChIP-seq datasets are normalised using Counts Per Million (CPM). RNA-seq that does not have stranded information is shown as a single track. For *Chlamydomonas reinhardtii* RNA polymerase 2 ChIP-seq is also displayed.
